# Phase separation of tunable biomolecular condensates predicted by an interacting particle model

**DOI:** 10.1101/2020.09.09.289876

**Authors:** Gorka Muñoz-Gil, Catalina Romero-Aristizabal, Nicolas Mateos, Lara Isabel de Llobet Cucalon, Miguel Beato, Maciej Lewenstein, Maria F. Garcia-Parajo, Juan A. Torreno-Pina

## Abstract

Phase separation is emerging as key principle in the spatiotemporal organization of living cells. Given its relevance in the regulation of numerous biological functions, including gene transcription and chromatin architecture, modeling biomolecular condensation is gaining interest. Yet, most models developed so far rely on specific descriptions and/or experimentally inaccessible properties. Here we propose a theoretical model, where phase separation is explained by means of interaction probabilities between particles. With minimum model requirements, particle condensates emerge above a critical interaction probability. We tested the model predictions with single molecule experiments of tunable transcription factor condensates in the nucleus of living cells. Phase separation, condensate sizes, diffusion behavior, and mobility parameters, quantified by data analysis and machine learning, are fully recapitulated by our model. Our combined theoretical and experimental approach provides a general framework to investigate the biophysical parameters controlling phase separation in living cells and in other soft matter-based interacting systems.

## Introduction

Activities performed by living cells are generally achieved through the compartmentalization of their multiple components in space and time. Although traditionally, cell compartments have been thought to be surrounded by membranes, recent evidence indicate that cells also organize membrane-less internal compartments through the physical process of liquid-liquid phase separation (LLPS)^1, 2, 3, 4^. LLPS creates transient chemically distinct compartments, also called biomolecular condensates, which might operate as versatile biochemical “hubs” inside the cell^1, 5^. Phase separation is particularly relevant in the cell nucleus, where the condensation of numerous proteins on chromatin have been shown to regulate gene transcription and chromatin architecture at multiple temporal and spatial scales^6, 7, 8^. Transcription factor (TF) condensates are proposed to regulate transcriptional initiation and amplify transcriptional output of expressed genes^5, 7, 9, 10, 11^. Yet, despite its prevalence and biological significance, a quantitative understanding of the biophysical parameters controlling transcription factor condensation in the living cell nucleus is largely missing.

Nuclear receptors are a family of TFs that have been widely studied as master regulators of gene transcription and genome topology in response to a tunable external stimulus: a steroid hormone^12, 13, 14^. Structurally, these TFs contain two intrinsically disordered regions: the N-terminal domain and the hinge (**Extended Data Fig. 1**); and two highly structured regions: the DNA-binding domain and the ligand-binding domain^15^. Ligand stimulation of several members of this family has been shown to trigger a LLPS, forming nuclear condensates with different transcriptional roles^16, 17, 18^. In the case of the progesterone receptor (PR), we observe similar condensate formation induced by progestin hormone (**Supplementary Movie 1**). Since it has been proposed that intrinsically disordered regions (IDR) are the main driving mechanism promoting LLPS in living cells^9, 19, 20, 21^, the nuclear receptor PR represents an ideal candidate to study inducible phase separation in a well-controlled and tunable experimental setting.

Theoretically, phase separation is usually associated to the heterogeneous mixing of two components, either by spinodal decomposition^22^, or nucleation^23^. In this context, entropy based models, as e.g. the Flory-Huggins model^24, 25^, have been commonly used to understand phase separated systems in biological scenarios^26^. Nevertheless, it is often very challenging to connect entropy with the actual physical interactions taking place between the particles of study. In the field of motile particles, phase separation has been extensively described as a collective behavior^27^. However, in the case of Brownian particles, phase separation can occur due to particle interactions, without the need of such collective phenomena, by introducing e.g. the possibility of binding^28, 29^, as is commonly considered in diffusion limited aggregation^30^. While all the previous methods show surprising insights on the phase separation phenomena, they often rely on very specific descriptions of the models, heavy numerical simulations, or experimentally inaccessible properties.

Here, we present a minimal particle-based interaction model that predicts phase separation with properties that are fully accessible to single molecule investigation in living cells. Analysis of single molecule trajectories combined with machine learning showed a bimodal mobility distribution and clearly distinct diffusion behavior of PR. Increasing the ligand concentration resulted in a sharp transition of spatiotemporal parameters and PR diffusion behavior above a critical ligand threshold, consistent with the emergence of condensates. Our minimal theoretical model: (1) predicts phase separation above a critical particle interaction threshold, (2) fully recapitulates the condensate properties obtained from our single molecule data, and (3) is independent of the underlying molecular mechanism by which the particles interact. TF condensation has been customary studied through ensemble or static measurements, mostly in *in-vitro* settings or in fixed cells. In contrast, the experiments and theoretical model presented here provide a general framework to investigate the dynamics of phase-separation in living cells at the single molecule level. Moreover, our approach can be further extended to a wide range of biological systems as well as other soft-matter based interacting systems.

## Results

### Single Particle Tracking of nuclear PRs in response to a tunable stimulus

Single Particle Tracking (SPT) has been widely used over the last decade to characterize the lateral mobility of many TFs and DNA binding proteins in the nucleus of living cells at the single molecule level^13, 31, 32, 33, 34^. We generated a stable MCF7 breast cancer cell line expressing a SNAP-GFP-PRB (PR isoform B)^35^. PR molecules were labeled with the SNAP-JaneliaFluor 549 (JF549) dye^36^ and their diffusion inside the nucleus was recorded under highly inclined illumination at a frame rate of 15ms (**Fig. 1a**). Individual JF549 localizations were reconnected to generate trajectories that report on the mobility of single PR molecules inside living nuclei. Individual trajectories were analyzed by computing the time averaged mean-square displacement (tMSD) and the angular distribution over consecutive steps (**Fig. 1b**)^32, 33^. The instantaneous diffusion coefficients for each trajectory were extracted by linear fitting of the 2^nd^-4^th^ points (D_2-4_) of the tMSD curve^37^; in this manner D_2-4_ histograms of hundreds of trajectories over different cells were generated (**Fig. 1b**, **Fig 1c)**. To investigate the PR lateral mobility in response to hormone, MCF7 cells were treated with different concentrations of the progesterone derivative R5020 (10^-12^ M to 10^-8^ M, 1h), or with EtOH as a control^38^. We observe two modes in the distribution of D_2-4_ values across different concentrations (**Fig. 1c**), similar to other proteins that interact with chromatin^31, 32^. Traditionally, these two components have been attributed to a chromatin-bound population (median D_2-4_ = 0.01 μm^2^/s) and to a freely diffusing population (median D_2-4_ = 0.5 μm^2^/s). Strikingly, instead of a gradual increase in the bound fraction of PRs that one would expect from a stochiometric occupancy of TFs to DNA binding sites with increasing ligand concentration, we found a sharp transition from freely to bound fraction taking place at a critical ligand concentration (10^-10^ M). This sharp transition in PR mobility suggests that LLPS might be regulating the interaction between PR and chromatin.

**Fig. 1:**
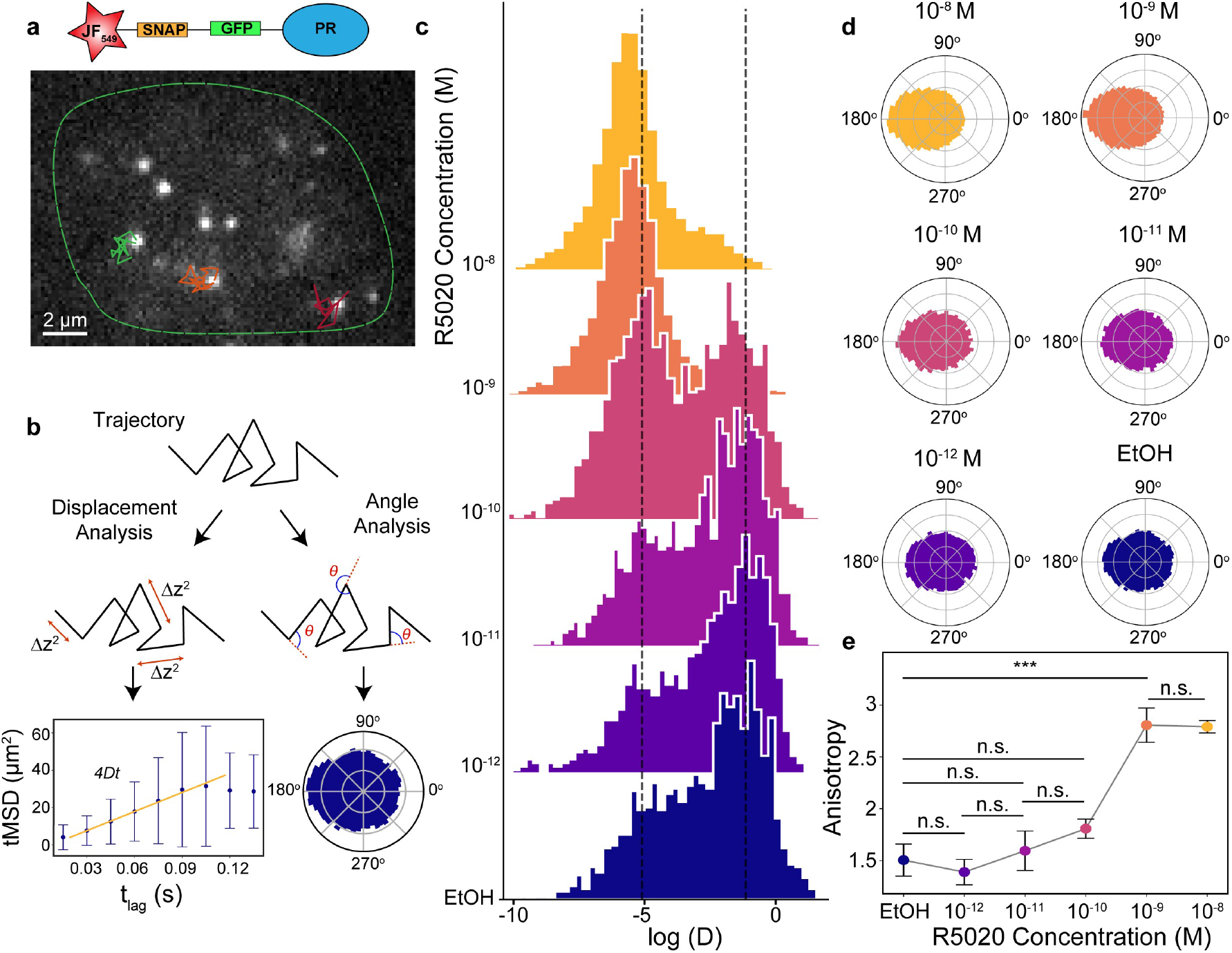
SPT analysis of the lateral diffusion of PR in living cells. **a**, Representative video frame of a SPT video. Individual PR molecules were visualized in the nucleus (green outline) of a MCF7 breast cancer cell line, under a highly inclined illumination at 15 ms frame rate. Diffraction-limited single molecule localizations are tracked in successive frames to generate individual trajectories (color lines). **b**, Schematic representation of the trajectory analysis. For each trajectory we extract the displacement between frames to generate individual tMSD plots as a function of the time lag and extract the diffusion coefficients (D_2-4_) for each trajectory (left, lower panel). In addition, we calculate the angles between successive steps to create polar histograms (right, lower panel). **c**, Distribution of the D_2-4_ (μm^2^/s) values of individual PR trajectories exposed to increasing R5020 concentrations for 1h. EtOH corresponds to the control condition, i.e., in the absence of ligand. Y axis corresponds to the frequency of events. Vertical dash lines indicate D_2-4_ values 0.0061 and 0.5 μm^2^/s. Data extracted from at least 1000 trajectories belonging to at least 8 cells from 3 independent experiments. **d**, Polar histograms of the angle between successive steps of diffusing PR under increasing R5020 concentrations. **e**, Anisotropy values as a function of R5020 concentration. One-way ANOVA test. n.s., not significant, ***, p-value<0.001.

We further computed the angle distribution for each individual trajectory (**Fig. 1d**). At hormone concentrations 10^-10^ M and below, diffusion is mainly isotropic and PR explores all angles with equal probability. In strong contrast, above the critical concentration of 10^-10^ M, the angle distributions become highly anisotropic with an increased occurrence of angles at 180°, i.e., higher probability for PR molecules to bounce back to their prior position. We computed the degree of anisotropy as the fold increase of angles occurring at 180^0^±30^0^ with respect to 0^0^±30^0^ (**Fig. 1e**)^32^. A sharp transition in anisotropy is retrieved above 10^-10^ M R5020 concentration, alike to that at which the D_2-4_ sharp transition takes place. We interpret this preferential backward movement as evidence of confinement, and a plausible indication of the bias in angles experienced by a particle inside a condensate when being constrained by the condensate boundaries. Altogether, our SPT results are consistent with a ligand-tunable and regulated LLPS process.

### Diffusion behavior of PR determined with machine learning

To identify the diffusion behavior of PR as function of hormone concentration, we relied on a recently developed machine learning (ML) analysis^39^. Using this algorithm we: (1) identify the theoretical model that best describes the diffusion behavior of individual PR trajectories, and (2) determine the corresponding anomalous exponent *α*, defined as the scaling factor when fitting the tMSD to a power-law ~ t^*α*^. Here, *α* = 1 corresponds to Brownian diffusion, *α* < 1 to anomalous sub-diffusion, and *α* > 1 to super-diffusion.

We first trained the algorithm with a set of simulated trajectories arising from various diffusion models related to many different experimental observations (see Methods). When applied to the experimental data, the vast majority of the trajectories were either classified as diffusing according to the annealed transit time model (ATTM)^40^, or exhibiting fractional Brownian motion (FBM)^41^. ATTM has been associated to the anomalous, non-ergodic and non-Gaussian motion of particles diffusing in a spatiotemporal heterogeneous medium, e.g. on cell membranes^42^. FBM has been described as an extension of Brownian motion with correlated noise, often associated to diffusion in viscoelastic media^43^. Note that, since the trajectories are normalized before entering the ML architecture (see Methods), the ML prediction is independent of the diffusion coefficient value.

For each hormone concentration, we computed the percentage of trajectories predicted as ATTM or FBM. At low ligand concentrations (<10^-10^ M R5020), 60% of the trajectories are classified as ATTM and 40% to the FBM model (**Fig. 2a**). Notably, a sharp change in the diffusion behavior occurs at >10^-10^ M R5020, with ~80% of the trajectories exhibiting FBM and ~20% ATTM (**Fig. 2a**). We further computed the D_2-4_ values of the trajectories assigned to each of the theoretical models and found that FBM trajectories display a much lower lateral mobility as compared to those assigned to ATTM (**Fig. 2b**). These findings indicate that the diffusion behavior of lower mobility trajectories may result from viscoelastic interactions between PR and chromatin within a condensate.

**Fig. 2:**
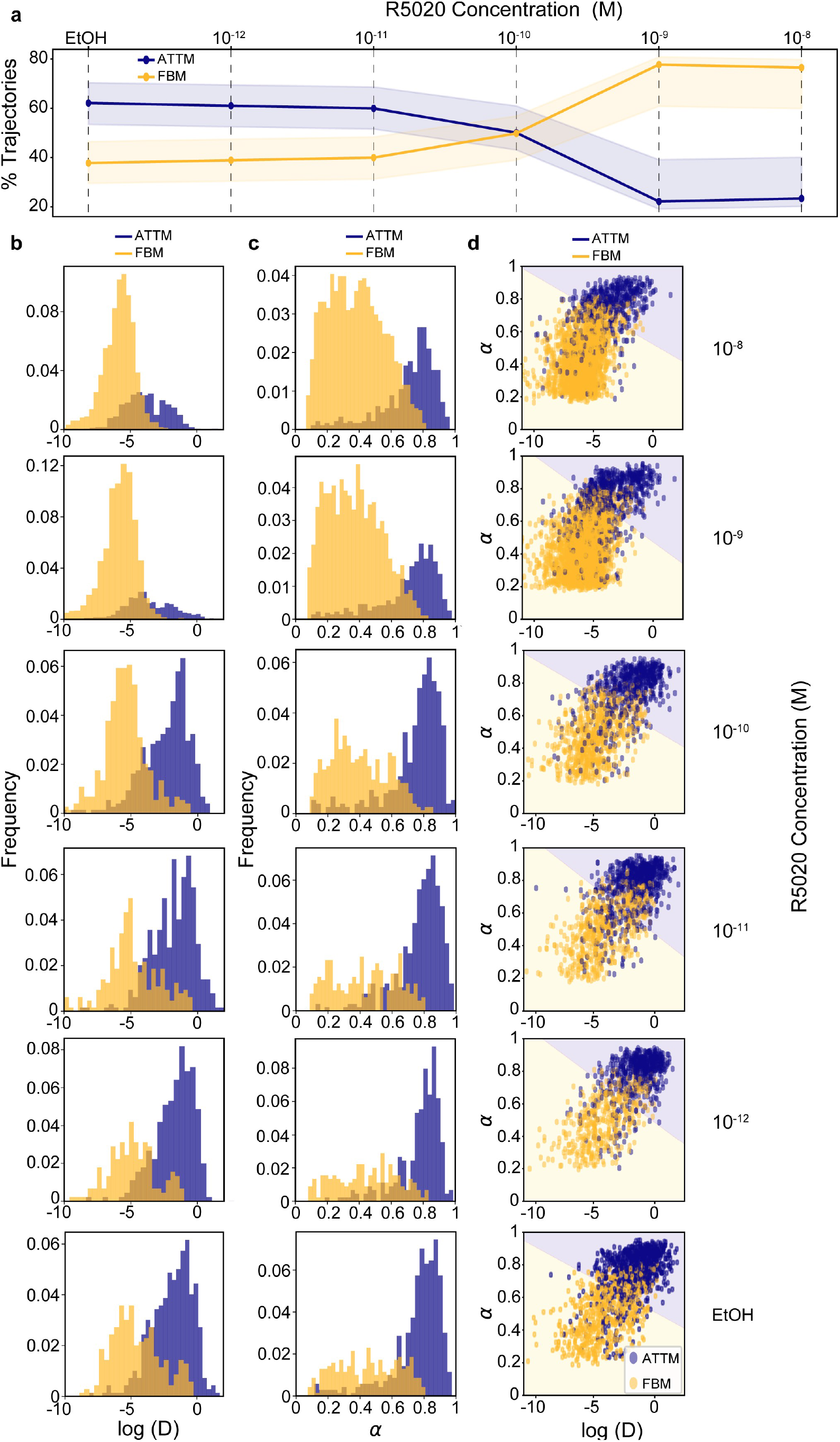
Machine learning analysis of individual PR trajectories in living cells. **a**, Percentage of trajectories associated to ATTM (blue) or FBM (yellow) by the ML algorithm as a function of ligand concentration. The shadow areas represent the error of the prediction, calculated by means of a confusion matrix (see Methods) **b**, D_2-4_ (μm^2^/s) distributions as a function of ligand concentration, with trajectories associated to ATTM (blue) and FBM (yellow) as identified by ML. **c**, Corresponding histogram of the ML predicted anomalous exponents. **d**, Scatter plot of the D_2-4_ vs. anomalous exponent for every trajectory. Background color represents the prediction of a SVM trained on the data.

Using a different ML architecture (see Methods), we further predicted the *α* values for each of the observed trajectories. FBM trajectories showed in average lower *α* values (~ 0.45) than ATTM trajectories (~ 0.75) (**Fig. 2c**). To assess the relationship between *D* and *α*, we generated scatterplots for different ligand concentrations (**Fig. 2d**). Strikingly, trajectories assigned to either ATTM or FBM form two differentiated clusters, that can be readily classified by a support vector machine (SVM). The background color of **Fig. 2d** shows the predictions of the SVM, demonstrating that *α* and *D* are sufficient to separate the lateral diffusion behavior of individual PRs as a function of ligand concentration. Overall, the ML analysis accurately separates two PR populations diffusing in markedly different media; and most importantly, it reflects a critical ligand concentration at which a transition from unbound (ATTM) to chromatin-bound (FBM) takes place.

### Nanometer-scale spatiotemporal mapping of PR in the nucleus

To further investigate the PR condensates from a single molecule perspective, we generated 2D spatiotemporal maps of the nanometer localization positions of individual PRs (using SNAP-GFP-PRB), as they dynamically explore the nuclear region with a ligand concentration of 10^-8^ M (see Methods)^37^. Hormone treatment led to the increased clustering of single PRs localizations in the nucleus (**Fig. 3a**). Moreover, we readily observed merging events in time, consistent with LLPS (**Fig. 3b**). These merging events were further confirmed by confocal video imaging at high temporal resolution (see Methods) of fully saturated GFP-labeled PR molecules (**Fig. 3c**). Condensates arising from individual localizations were detected using a cluster algorithm, as developed for super-resolution single molecule localization methods^44^ (see Methods). Treatment with hormone led to a significant increase of PR localizations contained within condensates (median value: 49%) as compared to EtOH treatment (median value: 22%) (**Fig. 3d**). The number of condensates per area increased significantly in hormone exposed cells (**Fig. 3e**) and the number of particles per condensate was also significantly larger upon hormone stimulation as compared to control (**Fig. 3f**). Since the 2D maps contain both temporal and spatial information, we further calculated the lateral mobility of particles belonging to individual condensates by reconnecting their positions over consecutive frames. Remarkably, PR trajectories within condensates reproduce the mobility (**Fig. 3g**), angle distribution (**Fig. 3h**), and FBM diffusion behavior (**Fig. 3i**) of the slow population retrieved by SPT shown in **Fig. 1** and **2**. Overall, our combined approach provides experimental evidence at the single molecule level for ligand-dependent PR phase separation and condensate formation in living cells.

**Fig. 3:**
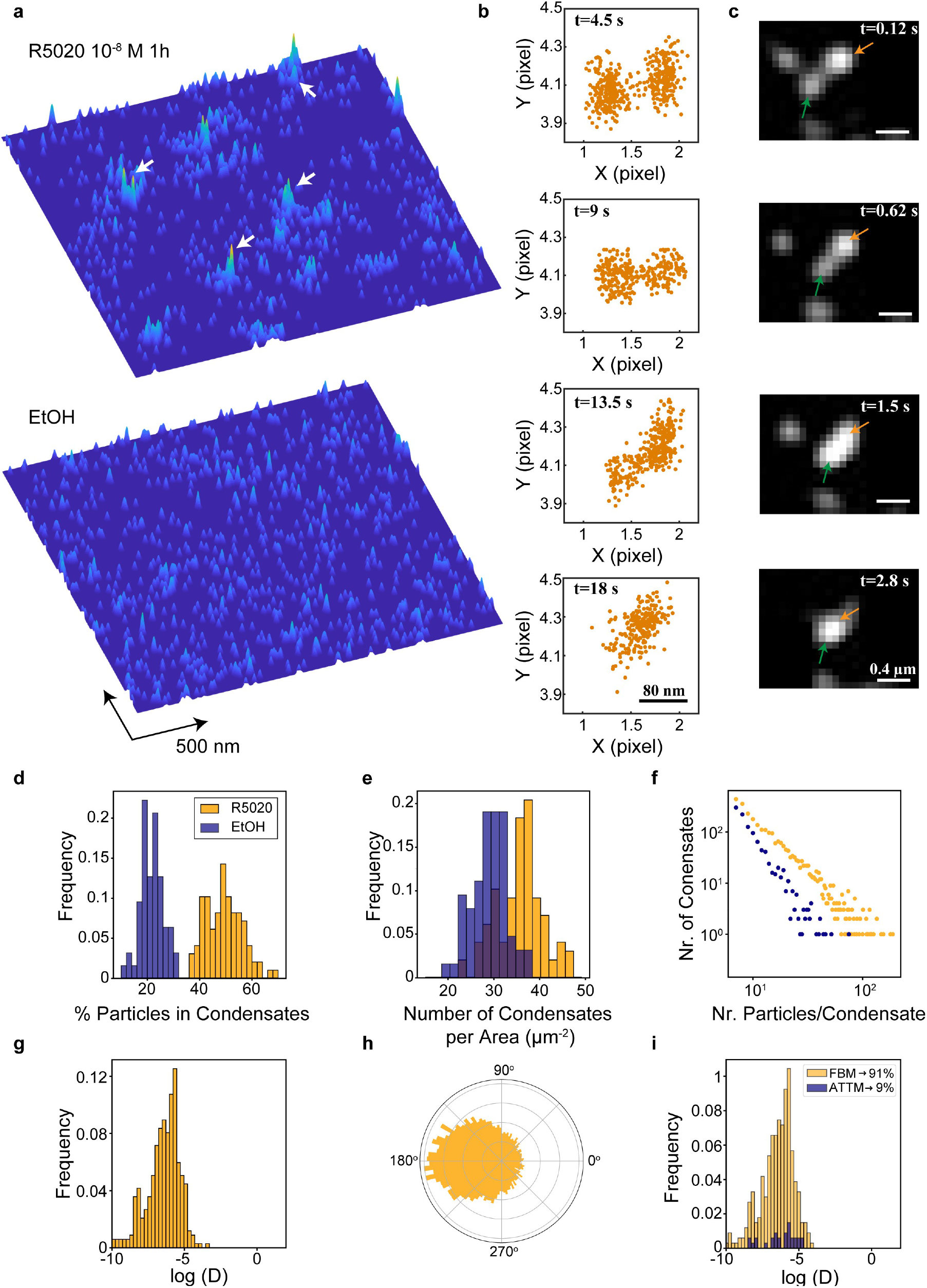
Nanometer-scale spatiotemporal mapping of PR in the nucleus. **a**, 2D localization maps of individual PRs over an area of 2.4 x 2.4 μm^2^ during 75 s, after 1h of ligand stimulation (upper panel) and control (lower panel). Each map contains 1000 localizations. White arrows indicate the presence of condensates. **b**, Snap shots of two different condensates as they merge over the indicated time windows. **c**, Merging events of different PR condensates visualized by confocal microscopy using GFP labeling conditions. **d**, Distribution of the percentage of particles within condensates. **e**, Distribution of the condensate density (i.e. number of condensates per unit area). **f**, Distribution of the number of particles per condensate. **g**, Distribution of the D_2-4_ values of PR trajectories inside condensates and corresponding **h**, Angle distribution between successive steps, and **i**, ML trajectory assignment to the diffusion behavior. Data from a minimum of 12 cells, in at least two independent experiments.

### Theoretical model based on particle interactions

To rationalize our results we developed a theoretical model in which particles-PR dimers in our case (see Methods), or other biological components in a general context-diffuse freely through the system, but also interact with each other in a non-trivial way. Indeed, we envisage that in a phase separation process particles most likely interact between each other with a certain binding probability, forming small condensates that practically immediately dissolve. Only when such binding probability is higher than a given critical value, condensates can grow, segregating themselves from their surrounding medium, and thus forming a new phase.

To model this process, we consider *N* particles with a space of action r that randomly diffuse in a system of size *S*. The model we consider is valid in any dimension with *r* and *S* having equal units. If the space of action of two particles overlap, they have a probability *P_c_* of binding. This mechanism thus favors the appearance of condensates, where *M* denotes the number of particles in a given condensate. Once formed, dissolving a condensate is less favorable than forming, in such a way that the probability of unbinding is 1 – *P*′_*c*_, with 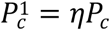 and 1 ≤ *η* ≤ 1/*P_c_*. With this, the probability that *n* particles escape from a condensate with *M* particles is given by:

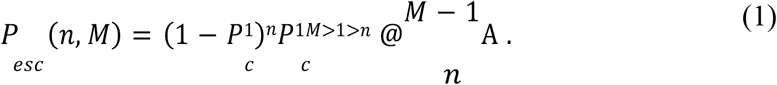

Using equation (1), we calculate the mean number of particles escaping from the condensate as 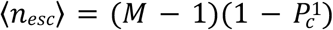 (**Fig. 4a**). Similarly, we calculate the mean number of particles being absorbed by a condensate of size *M* 〈*n_abs_*〉 (see Methods) (**Fig. 4b**). While 〈*n_esc_*〉 has the expected linear behavior, 〈*n_abs_*〉 grows non-linearly with *M* until reaching *M* = *N*/2, where the condensates become so large that the probability of finding new particles in the system starts to decrease, and so does 〈*n_abs_*〉. Notice that larger condensates have a higher probability of absorbing particles than smaller ones. This accounts in our model for classical mechanisms of droplet growth such as Ostwald ripening or diffusion limited growth^45^.

**Fig. 4:**
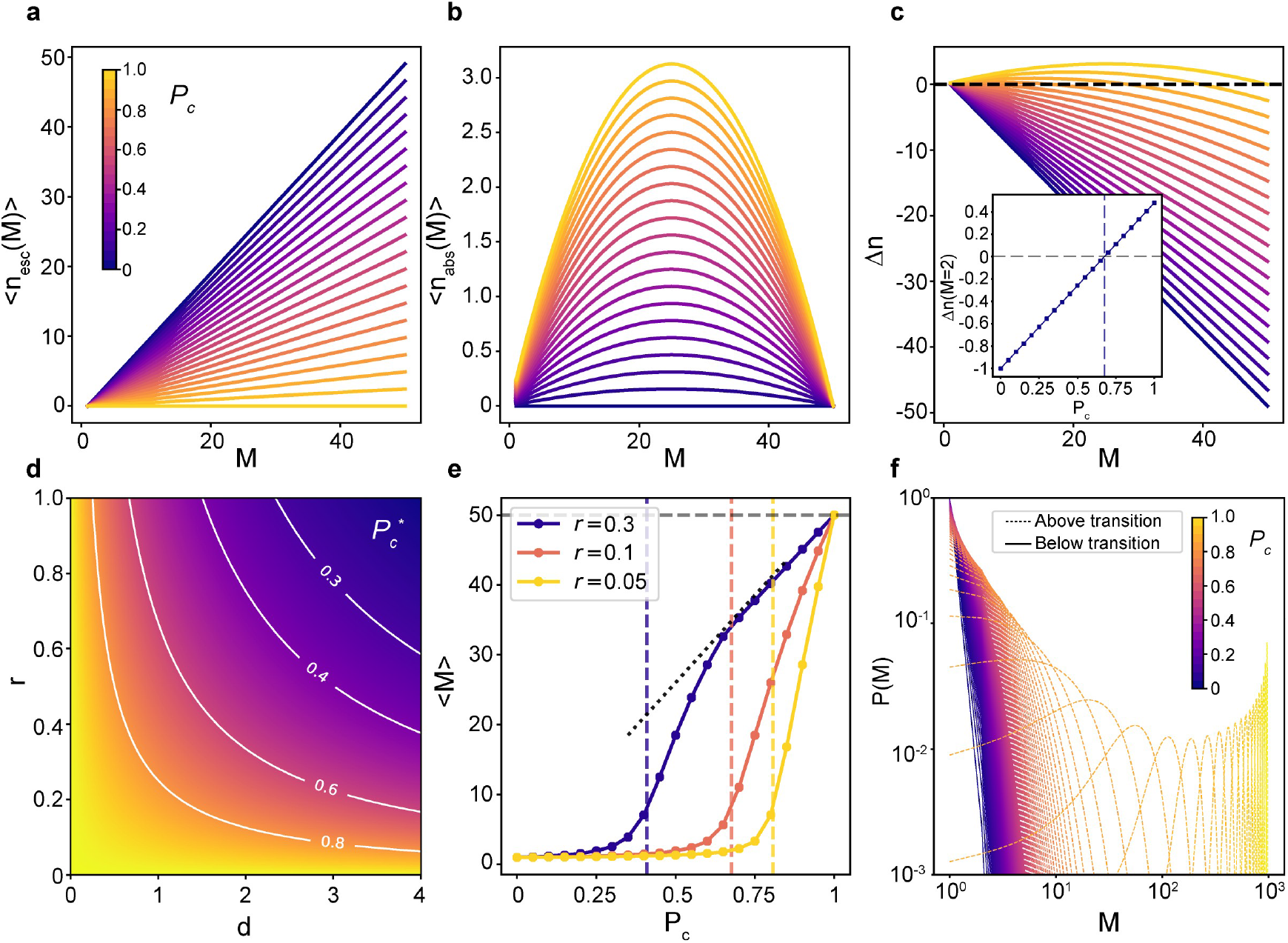
Particle-based interaction model predicting phase transition and condensate formation. Solutions are given for a system with *N* = 50, *S* = 10, *η* = 1 and *r* = 0.05, unless otherwise indicated. **a**, Mean number of particles escaping from a condensate of size *M*. **b**, Mean number of particles absorbed by a condensate of size *M*. **c**, Flux of particles Δ*n* for a condensate of size *M*. Inset: Value of the flux at *M* = 2 as a function of *P_c_*. The point Δ*n*(*M* = 2) = 0 gives the critical 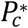. **d**, Critical 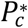 as a function of the density *d* = *N*/*S* and the space of action *r*. **e**, Average condensate size 〈*M*〉 as a function of *P_c_* for systems with different *r*. Vertical dashed lines represent the critical probability 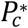. The dotted black line shows the prediction for the linear regime, valid for *r* = 0.3 at 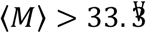. **f**, Condensate size distribution *P*(*M*) for various *P_c_* for a system *N* = 1000, *S* = (15*x*160 *nm*)^Z^, *η* = 1 and *r* = *π* (15 *nm*)^2^. Dashed (solid) lines show *P_c_* above (below) the critical 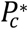.

Following our calculations, we determine the net flux of particles Δ*n* = (〈*n_abs_*〉 – 〈*n_esc_*〉 being on average absorbed or escaping from a condensate (**Fig. 4c**). In our system, condensate growth happens when Δ*n* > 0. Note, however, that only above a certain *P_c_*, the flux is positive for condensates of sizes *M* > 2, thus heralding the condensation. Such critical value 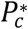 can thus be found by solving Δ*n*(*M* = 2) = 0 (inset of **Fig. 4c**). For systems containing a large number of particles such as proteins in a cell, 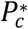 is well approximated by (see Methods for the exact solution):

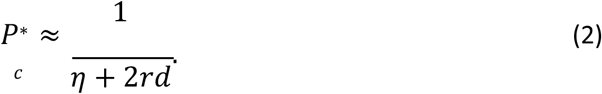

Since *d* = *N*/*S*, the critical 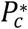 depends on the particles density and their corresponding space of interaction *r* (**Fig. 4d**). We then estimate the average size of condensates 〈*M*〉 for different *r*, corresponding to Δ*n*(< *M* >) = 0, as a function of *P_c_* (**Fig. 4e**). Importantly, condensates start rapidly emerging only above the critical binding probability 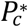 (vertical dashed lines in **Fig. 4e**), describing a new phase. In the hypothetical regime *rM* ≥ *S*, the flux scales linearly with *M* and we can obtain an analytical expression for 〈*M*〉 (see Methods and **Fig.4e**). Finally, we compute the distribution of condensate sizes *P*(*M*) by solving a system of recurrent equations, and by using the stochastic matrix theory^46^ (see Methods) (**Fig. 4f**). The size distributions are Gaussian with mean values that are strongly influenced by phase separation. Below the phase transition the means are always one with variance increasing with *P_c_*. Remarkably, above the phase transition, the means increase when increasing *P_c_* (**Fig. 4f**) and exactly coincide with 〈*M*〉. These distributions are relevant for the general case of soft-matter systems, but reflect an oversimplified scenario for living-cells, where we expect to find a limit to condensate size^45, 47^.

### Validation of the theoretical model to experimental single molecule data of PR in living cells

We begin our comparison by relating the parameters of the model to the experimental properties of our system. Since *P_c_* is defined as the binding probability between two particles, we consider that it is equivalent to the degree of PR activation by different ligand concentrations in our single molecule experiments. Unliganded PR is sequestered by a chaperone complex; only after its activation by hormone binding, PR is released and can interact with each other (see Methods). Biologically, this interaction could arise from weaker non-covalent binding strength between IDRs of PR due to π-stacking, charge interactions or non-polar residues among others^24^.

Condensates in living cells can only grow up to a given size^45, 47^, therefore we impose an upper limit to condensate size in our model: condensates rapidly dissolve after reaching *N_max_* particles, i.e. *P_c_* exponentially decreases for condensates with size above *N_max_*. With this consideration in mind, our theoretical model predicts the formation of condensates with sizes that follow a power law distribution above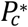 (**Fig. 5a**), similar to our experimental results in living cells (inset **Fig. 5a**). Moreover, the model further predicts a sharp decrease in the probability of finding free particles above 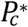(**Fig. 5b**), which fully agrees with the ML analysis of PR trajectories (inset **Fig. 5b**). Note that the peak close to the phase transition for *P*(1) and 〈*M*〉 is due to finite size effects. Finally, we perform a series of numerical simulations of the model. To account for the effect of condensation in the particle mobility, we assign a diffusion coefficient with mean *D*_T_ for freely diffusing particles, and mean *D_c_* for particles diffusing in a condensate, with *D_c_* < *D*_T_. For simplicity, we consider that all particles follow Brownian motion, as the theory is general for any diffusion model. After the system reaches its steady state, we generate histograms of the diffusion coefficients for all particles at various *P_c_*. We fully retrieve the bimodal distributions of our SPT experiments under different hormone concentration and notably, obtain a sharp change in diffusion above the critical *P_c_*(**Fig. 5c**). Thus, our model recapitulates the nature of a LLPS process making predictions that are amenable to single molecule investigation in living cells.

**Fig. 5:**
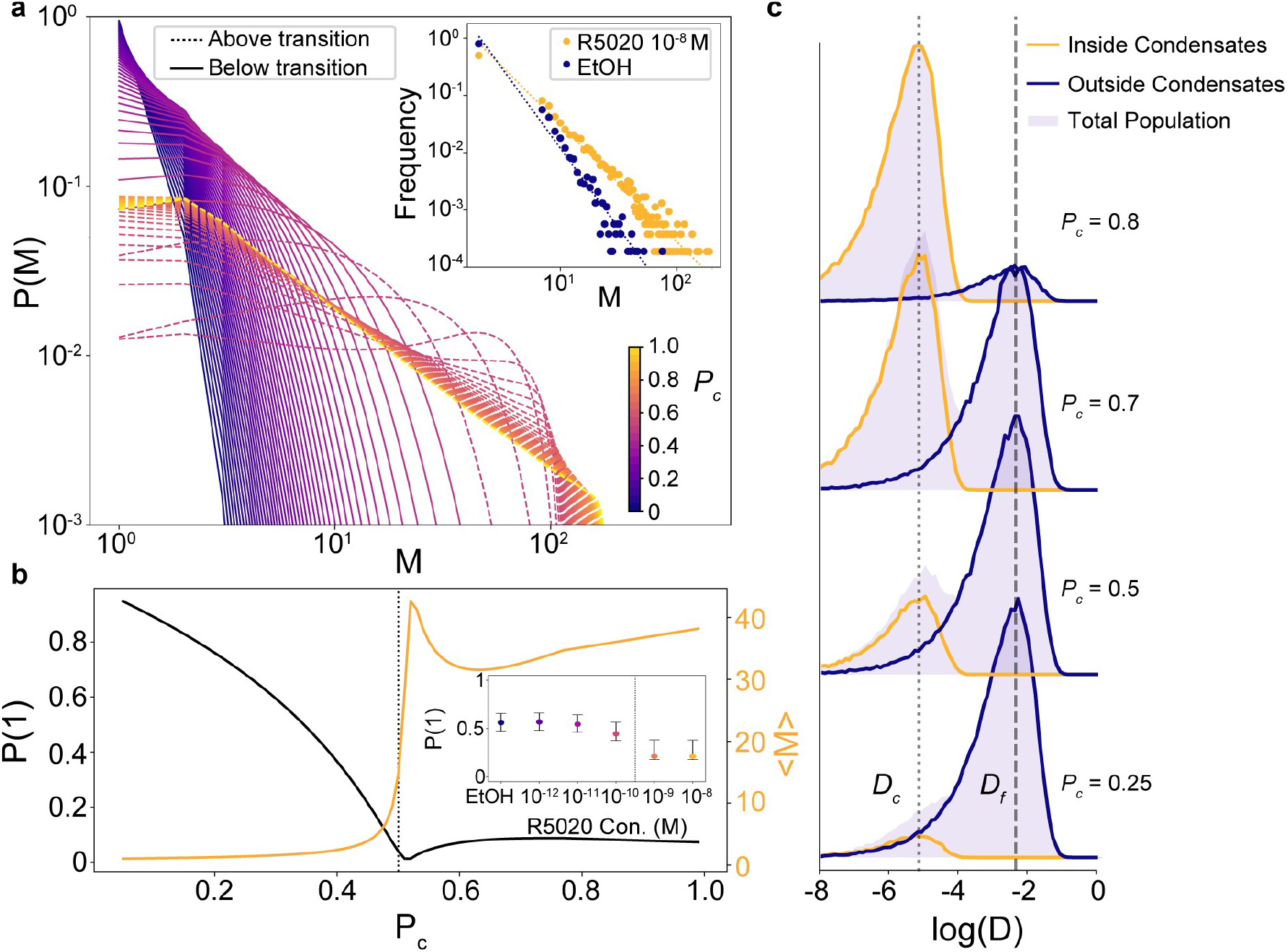
Validation of the theoretical model to experimental single molecule data of PR. **a**, Condensate size distribution *P*(*M*) for various *P_c_* for a system with *N* = 500, *r* = 0.1, *S* = 50, *η* = min (1.3, 1/*P_c_*) and maximum condensate size *N_max_* = 200. Dashed (solid) lines show *P_c_* above (below) the critical 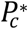. Inset: Experimental PR condensates size distribution of PR for control (blue) and hormone (yellow) at 10^-8^ M. Dashed lines show the corresponding power law fits. **b**, Black line: probability P(1) of finding free particles for the system in **a**, as a function of *P_c_*. Yellow line: normalized mean sizes of the condensate for the system in **a**. Dotted line: critical 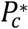 predicted by Eq. (2). Inset: Fraction of free PR molecules as function of ligand concentration, extracted from the ML analysis of experimental data on PR trajectories predicted to exhibit ATTM diffusion (see **Fig. 2a**). **c,** Distributions of D_2-4_ (μm^2^/s) values for numerical simulations on a system with *N* = 1000, *S* = (15*x*160 *nm*)^Z^, *η* = 1 and *r* = π (15 *nm*)^2^, for various values of *P_c_. D_c_*, diffusion of particles inside a condensate, *D*_T_, diffusion of free particles outside condensates.

## Discussion

Our experiments and theoretical model provide a better understanding of transcription factor phase separation at the single molecule level in the nucleus of living cells. We have introduced a general model which, with minimal considerations, fully reproduces the key features of an experimental system undergoing phase separation. While there exists a plethora of theoretical models able to describe such phenomenon^3^, they are either based on experimentally inaccessible properties, rely on numerical simulations, or are system dependent. In contrast, our model is based on the sole consideration that diffusing particles may interact one with another and will form condensates above a critical interaction probability. Importantly, this critical interaction probability depends on both, the concentration of the interacting particles and their corresponding individual interaction space, fully consistent with a phase separation process. Numerical simulations of diffusing particles inside the model-predicted condensates recapitulate our single molecule diffusion data. Our model further predicts that condensate sizes follow a Gaussian distribution with increasing mean values above the critical interaction point, applicable to a general context of soft-matter based interactions. Since it is highly improbable that such a steadily growing size distribution is found in living cells given their multiple regulatory mechanisms, we impose in our model that condensates increase up to a given size. With this additional assumption we fully recover the power-law distributions found in the nanoscale spatiotemporal maps obtained analyzing living cells.

Recent SPT experiments showed that TFs transiently bind to DNA with rather short binding times (in the seconds scale)^13, 14, 48^. We propose that condensate formation might increase the likelihood that individual PRs rebind within short timescales to their corresponding DNA binding region. Such a condensate environment will thus increase the effective time that a given DNA region is bound by TFs. This hypothesis is further substantiated by our experimental data, where FBM, traditionally associated to diffusion within viscoelastic media, was found to describe best PR low mobility diffusion. In conclusion, the unique combination of single molecule sensitive imaging techniques together with a general, minimal interaction model, as reported here, brings a substantial step forward in understanding the behavior of individual proteins within condensates. Overall, this work brings unique insight into phase separation in soft matter systems from both experimental and theoretical perspectives.

## ACKNOWLEDGEMENTS

We thank Gordon Hager for providing the pGFP-PRB plasmid and Luke Lavis for kindly providing the JF549 SNAP dye. We would like to thank the Advanced Light Microscopy Unit of the Center for Genomic Regulation (CRG, Barcelona) for their support. We thank G. Filion, R. Cortini, F. Campelo, M. A. García-March and C. Manzo for fruitful discussions. The research leading to these results has received funding from BIST-Ignite funding (PHASE-CHROM) (to C.R. and J.A.T.-P.), the European Commission H2020 Program under grant agreement ERC Adv788546 (NANO-MEMEC) (to M.F.G.-P.), ERC AdG NOQIA (to M.L.) and ERC Synergy Grant 609989 (to M.B.), Government of Spain (Severo Ochoa CEX2019-000910-S, FIS2017-89560-R (to M.F.G.-P.), JdC-IJCI-2017-33160 (to J.A.T.-P.), State Research Agency AEI (FIDEUA PID2019-106901GB-I00/10.13039 / 501100011033 (to M.L.), QuantumCAT _U16-011424 (to M.L.), cofunded by ERDF Operational Program of Catalonia 2014-2020, QUANTERA MAQS (funded by State Research Agency (AEI) PCI2019-111828-2 /321 10.13039/501100011033), (to M.L.)); Obra Social La Caixa (LCF-ICFO) (to G.M.-G.), Fundació CELLEX (Barcelona), Fundació Mir-Puig and the Generalitat de Catalunya through the CERCA program and AGAUR (Grants No. 2017 SGR 1341 to M.L. and No. 2017SGR1000 to M.F.G.-P.).

## AUTHOR CONTRIBUTIONS

J.A.T.-P., M.B., M.L. and M.F.G.-P. supervised the research. J.A.T.-P. and C.R.-A conceived the experiments. J.A.T.-P. performed the single molecule microscopy experiments and corresponding data analysis. C.R.-A. built the experimental system and performed the confocal microscopy. G.M.-G. and M.L. developed the theoretical model. G.M.-G. developed the machine learning architecture. N.M. performed data analysis. C.R.-A. and L.I.L.C. performed transfections and cell culturing experiments. All authors discussed the experiments and wrote the paper.

## METHODS

### Theoretical Model: Initial considerations

In classical physics there are three standard mechanisms for phase separation: spinodal decomposition, nucleation and diffusion-limited aggregation. Spinodal decomposition^1, 2, 3^ occurs for states that are unstable thermodynamically, i.e. correspond to, at least a local maximum of the free energy. Nucleation^4, 5, 6, 7^ is a mechanism, in which a new thermodynamic phase starts from a metastable state (a local minimum of the free energy). The time to nucleate can vary broadly depending how large is the energy barrier preventing nucleation. The mechanism most similar to the one considered in this paper is diffusion limited aggregation (DLA), whereby particles undergoing a random walk due to Brownian motion cluster together to form aggregates of such particles^8^. DLA theory focuses on spatial aspects and, for instance, the growth of the, so-called, Brownian trees. Similarly, it has been used to study the formation of aggregates of patchy particles^9^. Here we abstract from the spatial aspects of such systems, assuming that: 1) particles can bind with each other no matter their particular space orientation; 2) the clusters have circular shapes, as indicated by experiments. We consider that the binding probability *P_c_* accounts for the generalization of the former, as e.g. more complex spatial binding conditions can be described by a much lower binding probability.

### Theoretical Model: Absorbing probability

We present here in more detail how to calculate the probability *Pabs*(*n*, *M*) that a condensate of size *M* absorbs *n* new particles. First, the particle needs to be in the space of action of the condensate, *S_c_* ≈ *rM*. To account for this, we need to take into account the probability *P_s_* of a particle being in such space. Considering that the particle follows a diffusion model which explores space equiprobably in the long time limit, and also that *S* ≫ *r*, such that the previous condition is fulfilled, we have that *P_s_* ≈ *S_c_*/*S*. Thus, *Pabs*(*n*, *M*) can be written as

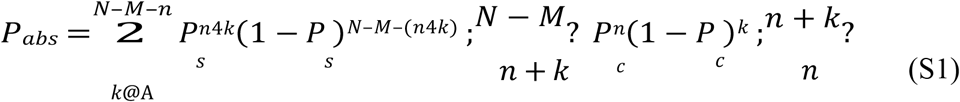

### Theoretical Model: Exact solution for critical *P_c_*

The critical binding probability 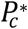 occurs when the flux is positive for any condensate size larger than two. Hence, one may find 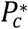 by solving the equation Δ*n*(*M* = 2) = 0, as shown in the inset of **Fig.4c.** While the flux has generally a quite complex form, it transforms to a much simpler form when considering *M* = 2, namely

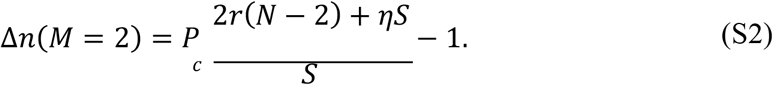

Equating Eq. (S2) to zero, we find that the critical probability is given by

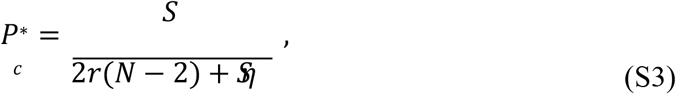

which in the case of *N* » 2, it is well approximated by 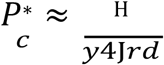.

### Theoretical Model: Linear regime

While the proposed theoretical model is fully solved analytically, due to the complexity of Eq. (S1) the resulting equations are quite involved. However, there exists a regime for which the system simplifies, and all the equations are linear with respect to the condensate size *M*. This happens, when the space of action is so large that the particle can indeed bind to any particle of the system. This effect may occur in cases of very long interactions, or in cases, when the particles’ size is of the same order of magnitude as the system. While these kind of systems are not very feasible, and in particular in the system of study here, the simplicity of the solutions and their connections with other physical systems, makes this limiting situation an interesting case of study.

We consider here the case for which *M* > *S*/*r*. This means that the space of action of the condensate *S_c_* ≈ *rM* > *S*, i.e., the condensate can bind to any particle of the system. Following the argument used to calculate Eq. (S1), we see here that the probability of a particle to be in the space of action of such condensate is *P_s_* = 1. This transforms *Pabs*(*n*, *M*) into

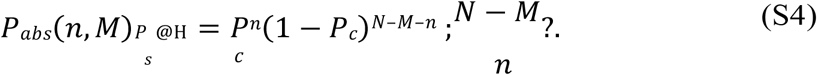

We can now calculate the average number of absorbed particles for such a condensate as

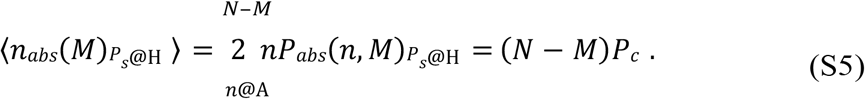

Now, using Eq.(1) from the main text, we can calculate the flux of particles as

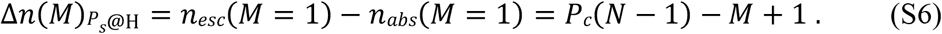

Finally, the mean size of the condensates can be calculated solving Δ*n*(*M*)_*p*_*s*@H__ = 0, to find

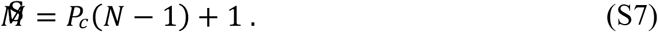

In **Fig. 4e** we plot such linear behaviour as dotted, for the case of *r* = 0.3 and *S* = 1, showing that the prediction matches for 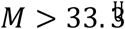, at which the system enters the linear regime.

### Theoretical Model: Condensate size distribution

To calculate the condensate size distribution used for **Fig. 4f** one needs to construct a system of recurrent equations. To understand this construction, let us consider a small system of three particles. Then, let us define *E_n,m_* = *P_esc_*(*n*, *M*) and *A_n,m_* = *P_abs_*(*n*, *M*). The probability of finding a condensate of size *M* = 1 is then given by

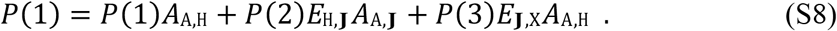

Similar equations are then constructed for *P*(2) and *P*(3). Such equations consider all possible ways of creating condensates of size *M* from all the rest of possible sizes ∈ [1, *N*]. Note that *E_m,m_* = 0 and that we always consider, both in simulations and in the theory, that escaping events take place before the absorbing ones.

In general, *P*(*M*) is given by

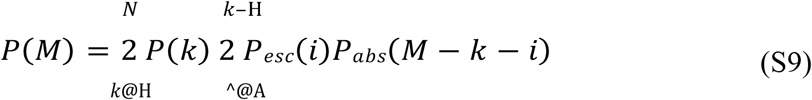

The above expression defines a set of *N* – 1 equations with *N* – 1 variables *P*(*M*), for *M* = 1,…, *N*. In order to solve such system, we consider the usual approach in stochastic matrix theory. A stochastic matrix (also called probability matrix) is a square matrix with non-negative real elements, each of them representing a probability^10^. They are often used to describe the evolution of a Markov chain. For instance, the set of equations generated by Eq. (S9) can be written in matrix form as

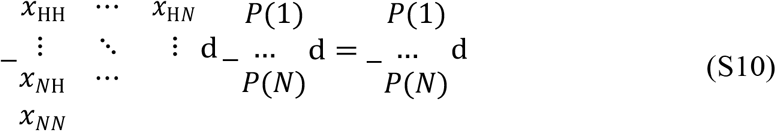

where *x*^_*j*_ are given by Eq. (S9). Note that the matrix *X*, the leftmost matrix in the previous equation, is a stochastic matrix. More precisely, it is a left stochastic matrix, as the sum of matrix elements in each of its columns is equal to one, i.e.

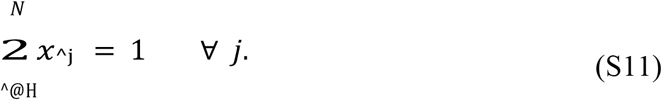

We can then use the stochastic matrix theory to solve the system of equations in question. For that, we give an initial ansatz for the vector of probabilities 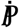. Then, we multiply this vector by the matrix *X*. The fact that *X* is a left stochastic matrix makes it such that the sum of the terms of 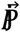 will always be equal to one, hence preserving the normalization of the condensate size probabilities. We consider for instance the ansatz 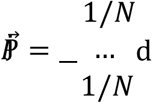. We then iterate this procedure until the convergence of 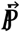.

### Machine learning algorithm

A schematic pipeline of the machine learning (ML) method used in this study is presented in **Extended Data Fig. 2**. The ML architecture is trained with a set of simulated trajectories. This tool allows to generate trajectories that are assigned to five different diffusion models. Moreover, trajectories with different anomalous exponents (0<*α*<1) can also be generated. The ML architecture can be trained separately to perform two different tasks: A) to classify the trajectories among a pool of different theoretical models; B) to regress the value of the anomalous exponent of each trajectory. Importantly, the training is done in a supervised way, i.e., we feed the trajectories to the machine, together with their corresponding labels (either the diffusion models for A, or the exponents for B). As architecture, we use a combination of gated recurrent units (GRU) and convolutional neuronal networks (CNN), merged with a contact layer made of fully connected neurons as depicted schematically in **Extended Data Fig. 2**. The GRU layers are able to learn long-term features, while the CNN are a good strategy to tackle short length correlations^11^. By combining the 2 approaches, we are able to characterize trajectories of only 10 data points in a robust manner.

### Machine Learning: Model classification

In order to classify the experimental trajectories according to a given diffusion model, the last layer of the network consists in a soft-max layer of *n* neurons, where *n* is the number of models considered. The labels are encoded in a vector of *n* elements, all equal to zero except the one encoding the model of the trajectory. The cost function to minimize is the Kullback-Leibler divergence which, for a set of trajectories 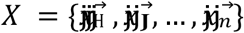 compares the output vector of the machine *f_m_*(*x*^) to the label vector 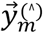 using

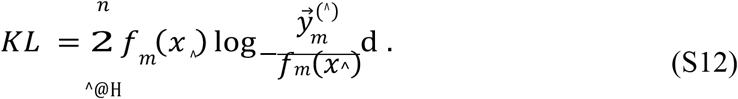

In order to faithfully characterize the set of experimental trajectories, we first train a model to classify among four diffusion models: continuous-time random walk (CTRW)^12^, fractional Brownian motion (FBM)^13^, annealed transient time motion (ATTM)^14^ and the scaled Brownian motion (SBM)^15^. For each model, we generate trajectories with anomalous exponent *α* ∈ [0.05,1], in intervals of 0.05. We create a balanced dataset with 1000 trajectories per model and exponent, which in total sum up to 72000 trajectories. We separate the dataset into two, a training set with 57600 trajectories and a test set with 14400. The latter will be used to calculate the accuracy of the model, i.e., to prevent the appearance of over-fitting. Note that the input size of the machine is fixed, which means that all the input trajectories should have the same size. As the experimental dataset has trajectories of varying size, from 10 to 1000 points, we solve such problem by cutting them to 20 frames long. This procedure ensures that most of the trajectories are considered, while the length is sufficiently big in order for the machine to have good accuracy. The trained model then has a micro-averaged F1-score of 0.733. When applied to the experimental dataset, 90 % of the trajectories were classified either as FBM or ATTM.

Since the vast majority of the trajectories were classified either as FBM or ATTM, we proceed to train the machine only with these two models. This allows to increase the accuracy of the ML classification for 20 frame long trajectories. In this case, the F1-score attained is of 0.822 (compared to 0.733). The confusion matrix for this classification is shown in **Extended Data Fig. 3a**. The results of the prediction on the experimental dataset are presented in the main text.

### Machine Learning: Anomalous exponent prediction

For the anomalous exponent prediction, the output of the machine is a continuous value. Hence, the last layer of the neural network is a single neuron with a rectifier activation function (RELU). The loss function in this problem is the mean absolute error,

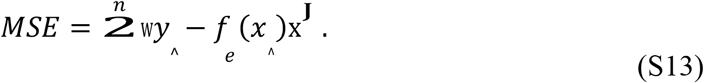

where *y*^ is the label corresponding the trajectory *x*^, and *f_e_*(*x*^) is the network prediction. The sum is done over the set of trajectories in the training dataset.

In order to infer the anomalous exponent for each individual trajectory, we used a simpler version of the neural network, containing 2 GRU layers of 100 and 50 neurons each, whose output enters 2 fully connected layers of 64 neurons and sigmoid activation functions. The last layer contains a single neuron with RELU activation function. Between each fully connected layer, we proceeded with a 25% dropout. This network shows a mean absolute error of 0.229 for trajectories of just 20 points. Note also that the predictions of the network are biased to increase the exponent, as shown in **Extended Data Fig. 3b**.

### Molecular mechanisms of progesterone receptor activation

Before the progesterone receptor (PR) is exposed to ligand, the PR is found both in cytoplasm and the nucleoplasm, where is tightly bound to chaperone complexes. These heat-shock proteins stabilize the extensive N-terminal disorder region to avoid proteolysis, which is intrinsically induced by disordered regions. The fact that the PR is sequestered by chaperones makes improbable its interaction with other PR molecules when the ligand is absent. Upon ligand binding, PR dissociates from chaperones, undergoes extensive post-translational modifications in its N-terminal region, dimerizes and migrates to the cell nucleus, where it binds palindromic DNA sequences similar to those bound by the glucocorticoid and the androgen receptors^16^. The fact that the binding site for PR is an inverted repeat, strongly suggests that the dimer of PR (as oppose to the monomer) can be considered as the prevalent functional unit. It has also been proposed that the active form of glucocorticoid receptor is a tetramer^17^, but whether this is the case for PR is unknown. In this manuscript we consider the dimers of PR as the functional unit for the transcriptional response. Thus, when we refer to “a particle” in the model, it corresponds molecularly to a PR dimer that has been translocated to the nucleus and it has the signature of post-translational modifications induced by hormone. We can then assume that by controlling the hormone concentration we are fine-tuning the volume fraction of active PR dimers in the cell.

### Plasmids

The original pGFP-PRB was a gift from Gordon Hager (National Cancer Institute, NIH, Bethesda, USA). This plasmid expresses the PR isoform-B under a tetracycline controllable promoter (TetOff system, Clontech). To perform the SPT experiments, a SNAP tag was introduced at the N-terminal to the GFP, using Gibson cloning (pSNAP-GFP-PRB). A Puromycin resistance plasmid (pPUR, Clontech, Cat No. 631601) was used as a selection marker. All plasmids were linearized with ScaI before electroporation.

### Cell culture and electroporation

MCF7 Tet-off cells (Clontech, Cat No. 631154) were grown on Dulbecco Modified Eagle Medium (DMEM) high-glucose media supplemented with 10% Tet-free Fetal Bovine Serum, 2mM L-glutamine, 1 mM sodium pyruvate, 100 U mL-1 penicillin and 100 μg mL-1 streptomycin. The cells were cultured at 37 °C in a CO2/air (5%/95%) incubator. Cells were electroporated simultaneously with the pSNAP-GFP-PRB and the pPUR, using a 10 to 1 ratio respectively. Electroporation was performed using the Amaxa Cell Line Nucleofector Kit V (Lonza) using the P-20 program, following manufacturer’s instructions. After one week, cells were selected under 0.6 ug/ml Puromycin, to enrich for electroporated cells, and then sorted in single cell wells using GFP as a marker, in order to generate a stable cell line.

### Hormone stimulation and SNAP labeling

Two days before the microscopy, 200 thousand cells were seeded in 35 mm glass bottom dishes. 16h before hormone stimulation, cells were washed with Phosphate-buffered saline solution, to eliminate traces of phenol red, and then changed to white DMEM media supplemented with 10% charcoal-treated FBS Serum, 2mM L-glutamine, 1 mM sodium pyruvate, 100 U mL-1 penicillin and 100 μg mL-1 streptomycin; from now on abbreviated as “charcoalized white DMEM”. The Janelia Fluor^®^549 dye coupled to the SNAP substrate was kindly provided by Luke Lavis (Janelia Farm, Ashburn, Virginia, USA). Cells were incubated with 10 nM for SPT and 100 nM for 2D spatiotemporal maps of the SNAP JF-549 dye in charcoalized white DMEM for 30 min at 37°C. Subsequently the cells were washed three times with PBS, and then placed back in the incubator in charcoalized white DMEM for a 1 h washout at 37°C. After the JF549 SNAP labeling, hormone stimulation was done using R5020 (Promegestone) solubilized in ethanol, or control conditions with this solvent. To study the response to different concentrations of hormone, a series of dilutions were made freshly before the microscopy acquisition.

### Experimental set-up for SPT and 2D spatiotemporal maps

Single particle tracking and 2D spatiotemporal maps imaging were performed in a Nikon N-STORM 4.0 microscope system for localization-based super-resolution microscopy, equipped with a TIRF 100x, 1.49 numerical aperture objective (Nikon, CFI SR HP Apochromat TIRF 100XC Oil). The sample was illuminated by a continuous 561 nm laser line with a power of 30 mW before the objective in HILO-configuration. The emission fluorescence of the JF549 was collected through the objective and projected into an EM-CCD Andor Ixon Ultra Camera at a framerate of 15 ms. The pixel size of the camera is 160 nm. During imaging, the temperature was kept at 37°C by an incubation chamber.

### Confocal imaging

GFP confocal line scanning microscopy was performed in a Leica TCS SP5 II CW-STED microscope using a 63x Oil Numerical Aperture 1.4 objective (Leica HC PL APO 63x/1.40 Oil CS), using a multiline Argon laser at 488 nm for excitation. The emission fluorescence was detected with a Hybrid detector (Leica HyD) in photon counting mode, using a 500–550 nm filtering. The sample was kept at 37°C with 5% CO2 by an incubation chamber. For **Fig. 3c** images of 256 x 256 pixels were acquired with pixel size of 80nm and dwell time of 9μs. Scanning was performed at 100Hz, acquiring consecutive frames every 125ms. For the **Supplementary Video 1** images of 322 x 200 pixels were acquired with a pixel size of 160nm. Each frame in the movie has a total integration time of 15 sec, and corresponds to the sum intensity projection from 100 images taken consecutively every 150ms, scanning at 700Hz.

### Generation of SPT trajectories

The nuclear region was segmented in the GFP channel intensity using Fiji. Individual tracks inside the nuclear region were analyzed using Trackmate^18^. Particle detection was performed with a Difference of Gaussians, with an expected diameter of 0.6 μm and sub-pixel localization. Detected particles were first filtered based on the Signal to Noise Ratio of the input image and then based on quality score. The particles retained were then linked using a simple Linear Assignment Problem (LAP) tracker, with a 1μm linking distance, 1μm gap-closing max distance and gap closing of two frames. Only tracks with more than 10 frames were considered for the analysis.

### Generation of 2D spatiotemporal maps

The total single molecule localizations of JF549 labeled PR molecules were detected by a custom Matlab Software over 5000 frames (75 s) and projected into one single frame. Condensates were detected by applying a Density-Based Spatial Clustering of Applications with Noise (DB-SCAN)^19^ over the entire frame with a threshold of 48 nm of interparticle distance and condensates containing a minimum number of particles of 5.

### Time-ensemble Mean Square Displacement and Diffusion Coefficients

Given trajectory whose two dimensional position (*x*, *y*) is sampled at *T* discrete, regular time steps *t*^, its time averaged mean-square displacement (tMSD) is calculated using^20^:

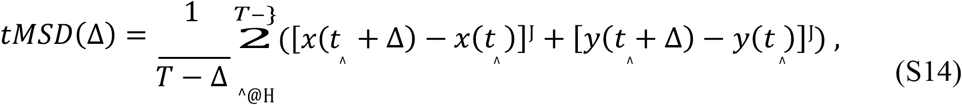

where Δ is usually referred as the time lag. Even in the presence of anomalous diffusion, at short times the MSD is well represented by

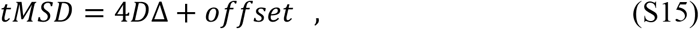

where *D* is the instantaneous diffusion coefficient. To extract it, we fit the tMSD between Δ = 2 to Δ = 4 and redefine it, as presented in main text, as *D*_J–ç_.

### Determination of the turning angles

For a given time, *t*, and a time between frames, *δt*, we define the turning angle, *θ_t_*, between consecutive trajectory segments, 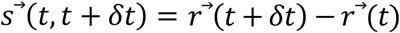, as follows^21^:

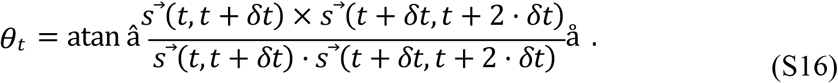

For our calculations, we consider the particle positions to be in 3D with the *z* component equal to zero. Using the above expression, the angles are defined between 0° and 360°.

### Determination of the turning angle anisotropy

To calculate the anisotropy of the turning angles, the fold change between the number of angles from 180°±30° and 0°±30° was extracted^22^.

## EXTENDED DATA FIGURES & FIGURE LEGENDS

**Extended Data Fig. 1:**
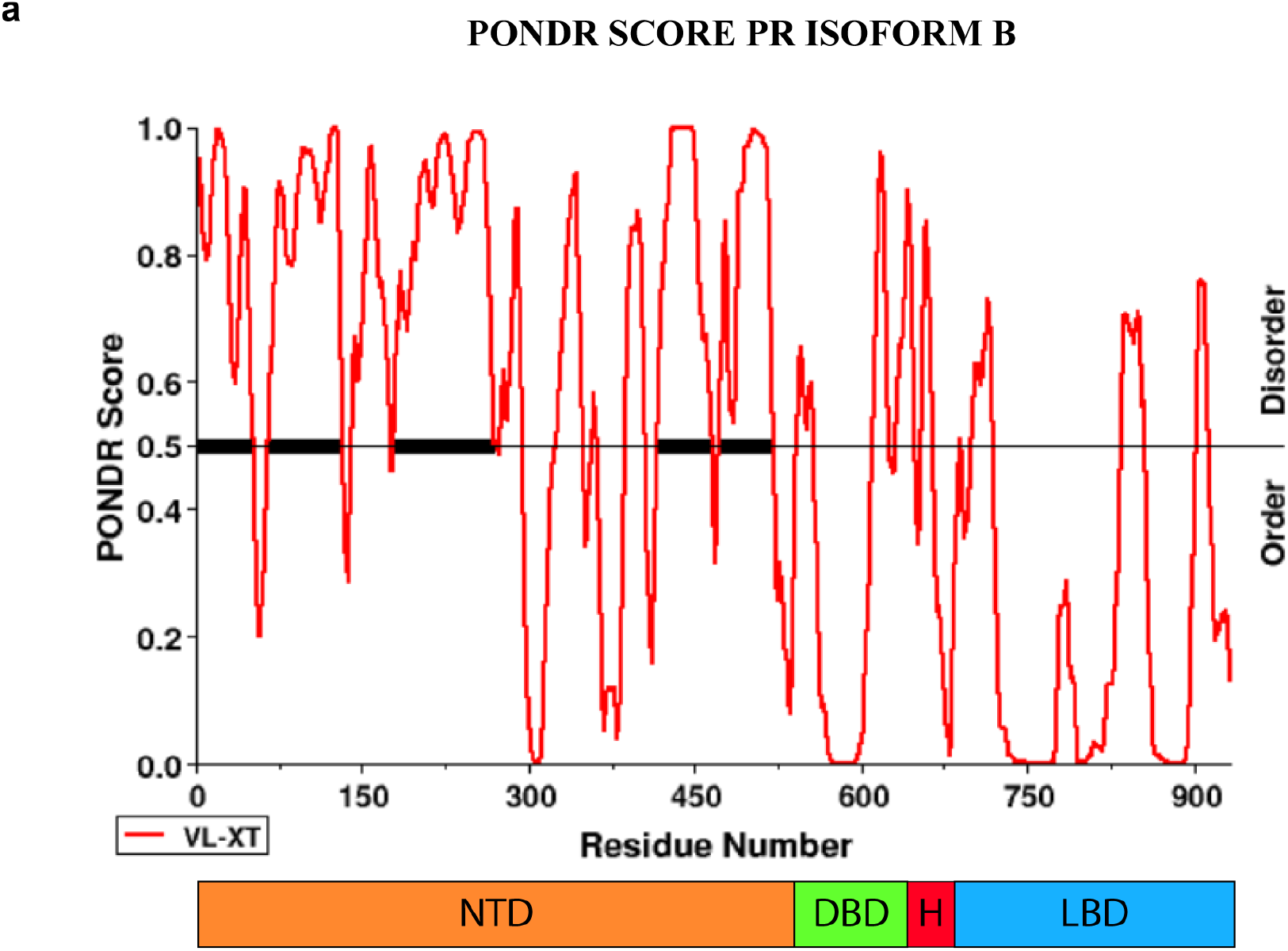
PONDR score of PR-B. Prediction of Natural Disordered Regions (PONDR score) of PR-B generated at www.pondr.com. Note the different regions of PR-B denoted as N-terminal domain (NTD), DNA-binding domain (DBD), the Hinge (H) and the ligand binding domain (LBD). The NTD is highly disordered (PONDR score > 0.5).

**Extended Data Fig. 2:**
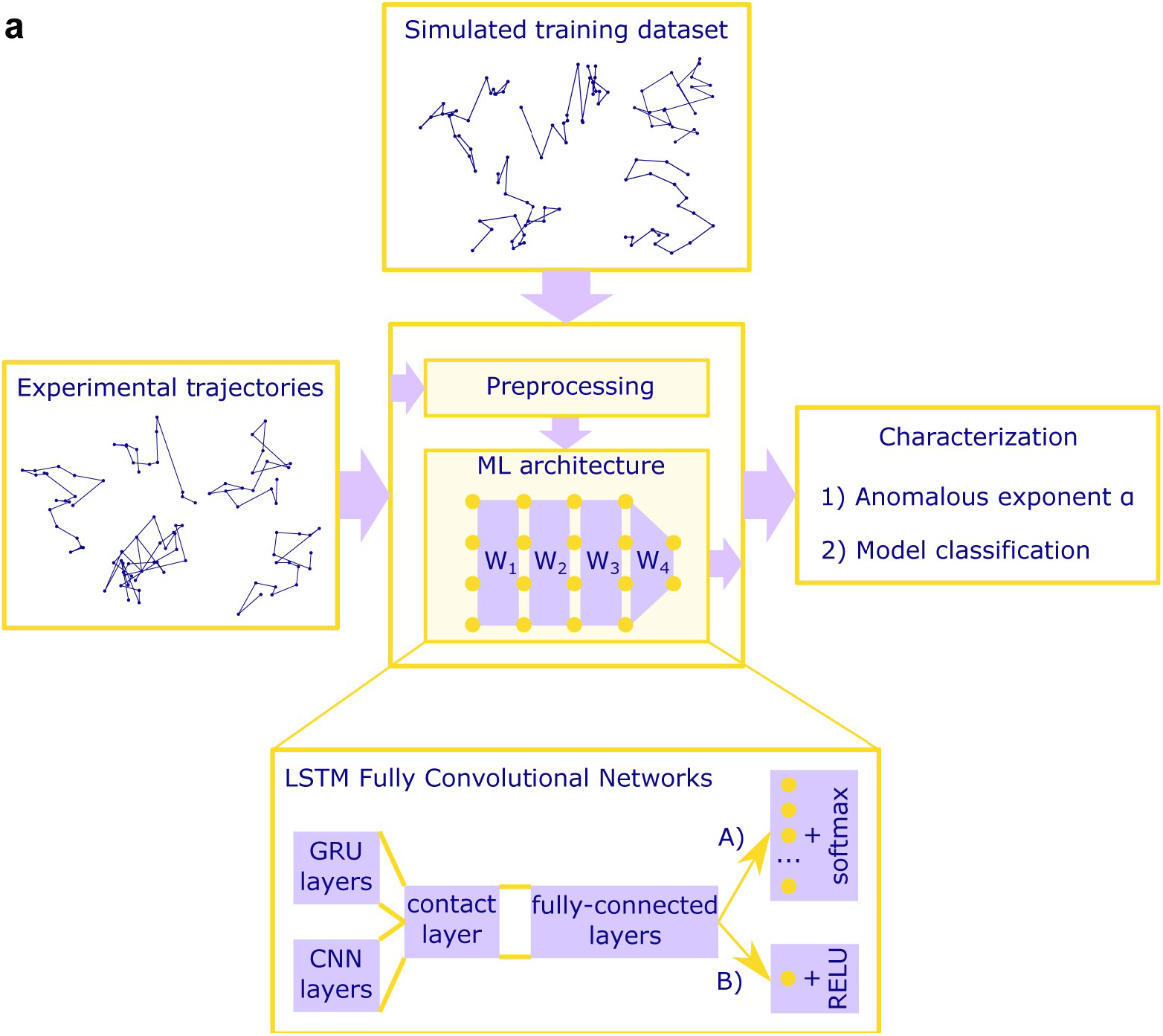
Scheme of the machine learning procedure. The machine learning (ML) architecture is trained with a dataset consisting on simulated trajectories. Once the training is complete, the machine can assign to every experimental trajectory an anomalous exponent and a diffusion model.

**Extended Data Fig. 3:**
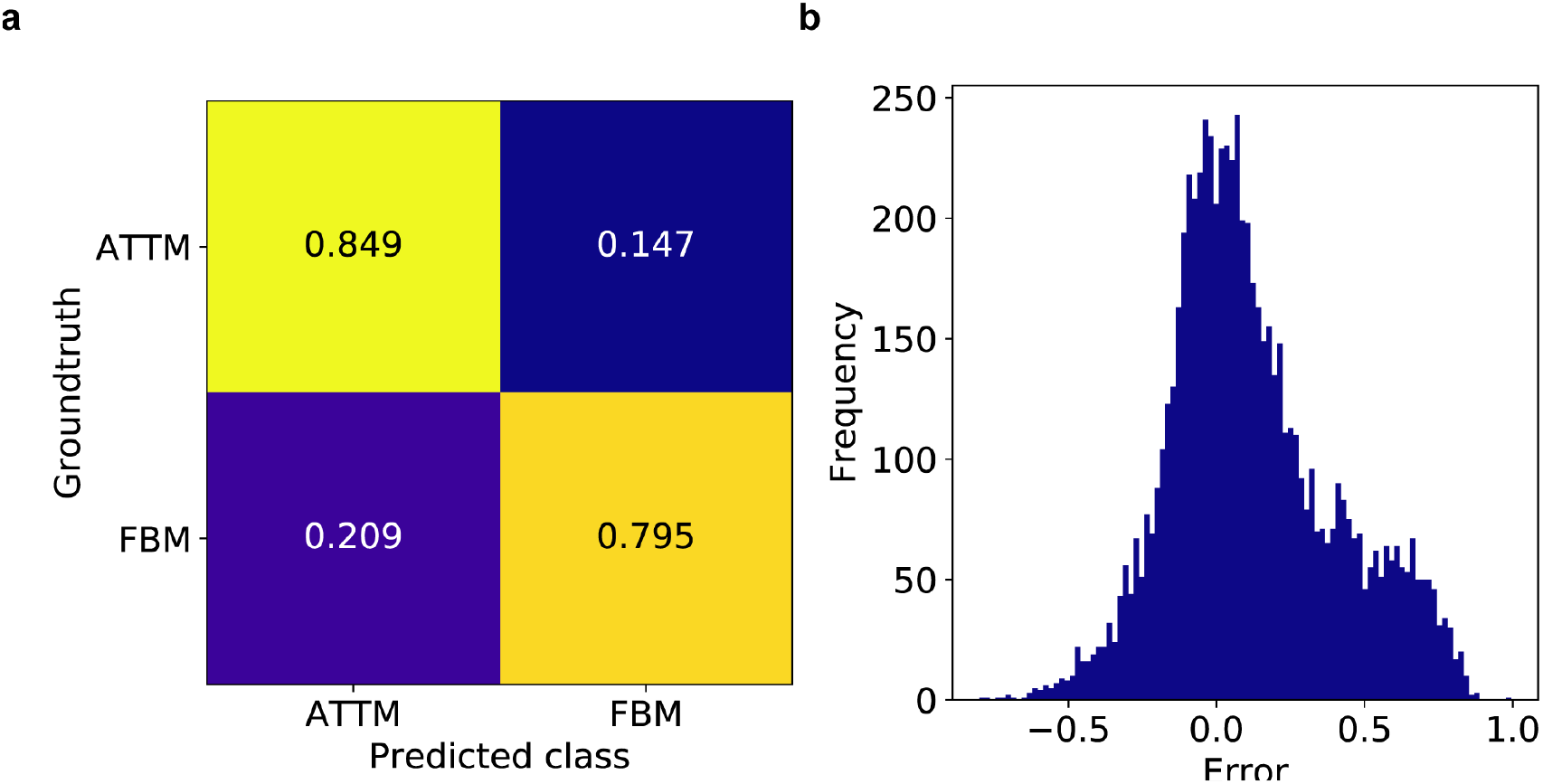
Error in the machine learning analysis. **a**, Confusion matrix for the long shot term memory neural network (LSTM) used for model classification. **b**, Prediction error for the gated recurrents units (GRU) network in the anomalous exponent prediction. In order to avoid overfitting, results were obtained using 7200 trajectories with T=20 frames in both cases.

**Supplementary Video 1:**
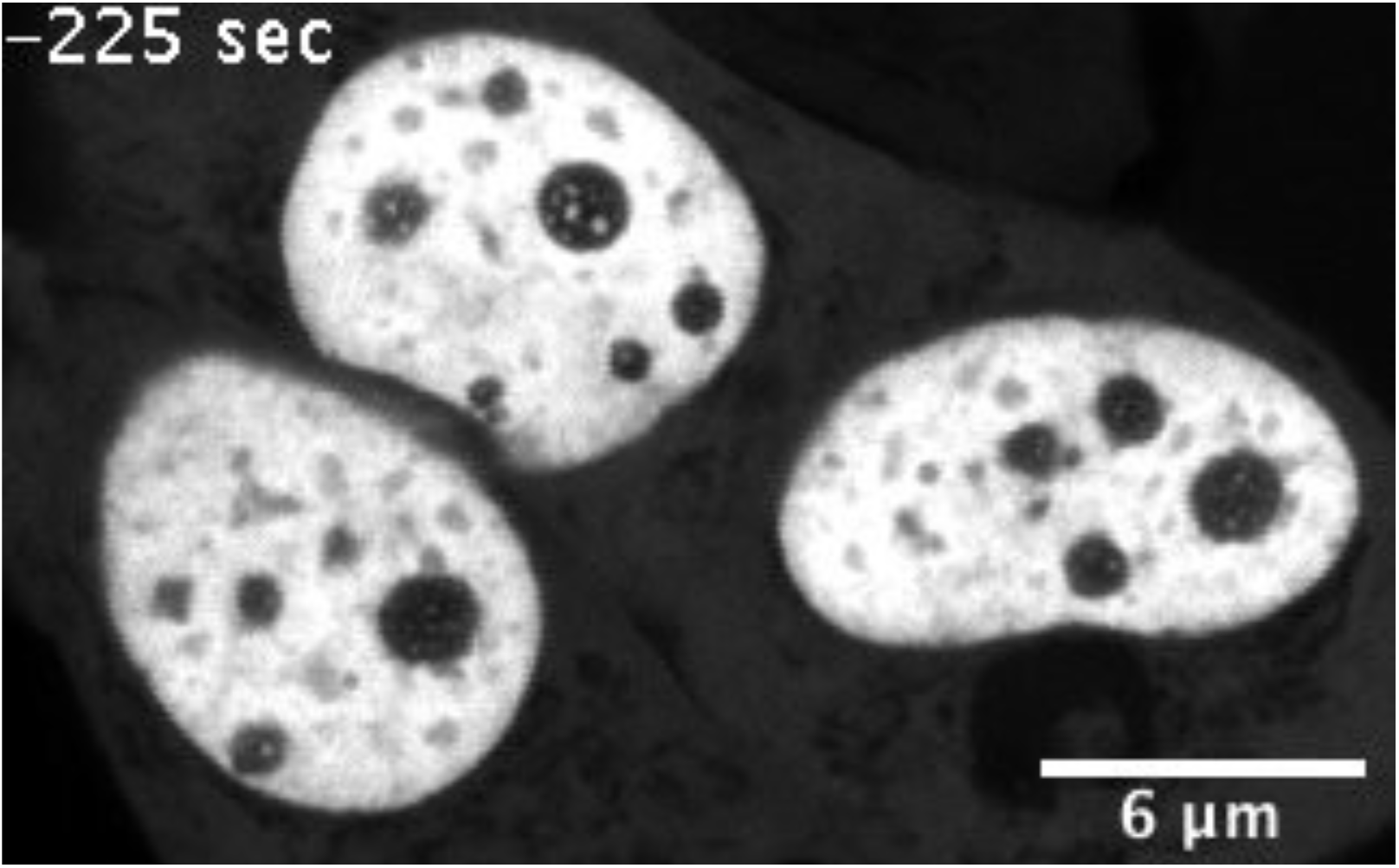
Time lapse of inducible PR nuclear condensates. MCF7 cell-line expressing GFP-PRB before and after hormone stimulation. Before treatment with hormone the fluorescent signal of the GFP-PRB is homogeneous across the nucleoplasm. After hormone addition (R5020 10^-8^ M, black frames) the fluorescent signal distributes into condensates within 5 minutes of hormone exposure. Each frame has a total integration time of 15 sec (see Methods).

## References

1. Banani, S.F., Lee, H.O., Hyman, A.A. & Rosen, M.K. Biomolecular condensates: organizers of cellular biochemistry. Nat Rev Mol Cell Biol 18, 285–298 (2017).

2. Brangwynne, C.P. et al. Germline P granules are liquid droplets that localize by controlled dissolution/condensation. Science 324, 1729–1732 (2009).

3. Hyman, A.A., Weber, C.A. & Julicher, F. Liquid-liquid phase separation in biology. Annu Rev Cell Dev Biol 30, 39–58 (2014).

4. Shin, Y. & Brangwynne, C.P. Liquid phase condensation in cell physiology and disease. Science 357(2017).

5. Chong, S. et al. Imaging dynamic and selective low-complexity domain interactions that control gene transcription. Science 361(2018).

6. Cai, D. et al. Phase separation of YAP reorganizes genome topology for long-term YAP target gene expression. Nat Cell Biol 21, 1578–1589 (2019).

7. Guo, Y.E. et al. Pol II phosphorylation regulates a switch between transcriptional and splicing condensates. Nature 572, 543–548 (2019).

8. Larson, A.G. et al. Liquid droplet formation by HP1alpha suggests a role for phase separation in heterochromatin. Nature 547, 236–240 (2017).

9. Boija, A. et al. Transcription Factors Activate Genes through the Phase-Separation Capacity of Their Activation Domains. Cell 175, 1842–1855 e1816 (2018).

10. Sabari, B.R. et al. Coactivator condensation at super-enhancers links phase separation and gene control. Science 361, 3958 (2018).

11. Shrinivas, K. et al. Enhancer features that drive formation of transcriptional condensates. Mol Cell 75, 549–561 e547 (2019).

12. Le Dily, F. et al. Hormone-control regions mediate steroid receptor-dependent genome organization. Genome Res 29, 29–39 (2019).

13. Paakinaho, V. et al. Single-molecule analysis of steroid receptor and cofactor action in living cells. Nat Commun 8, 15896 (2017).

14. Stavreva, D.A. et al. Transcriptional Bursting and Co-bursting Regulation by Steroid Hormone Release Pattern and Transcription Factor Mobility. Mol Cell 75, 1161–1177 e1111 (2019).

15. Hill, K.K., Roemer, S.C., Churchill, M.E. & Edwards, D.P. Structural and functional analysis of domains of the progesterone receptor. Mol Cell Endocrinol 348, 418–429 (2012).

16. Bouchard, J.J. et al. Cancer Mutations of the Tumor Suppressor SPOP Disrupt the Formation of Active, Phase-Separated Compartments. Mol Cell 72, 19–36 e18 (2018).

17. Nair, S.J. et al. Phase separation of ligand-activated enhancers licenses cooperative chromosomal enhancer assembly. Nat Struct Mol Biol 26, 193–203 (2019).

18. Stortz, M., Pecci, A., Presman, D.M. & Levi, V. Unraveling the molecular interactions involved in phase separation of glucocorticoid receptor. BMC Biol 18, 59 (2020).

19. Brady, J.P. et al. Structural and hydrodynamic properties of an intrinsically disordered region of a germ cell-specific protein on phase separation. Proc Natl Acad Sci U S A 114, E8194–E8203 (2017).

20. Elbaum-Garfinkle, S. et al. The disordered P granule protein LAF-1 drives phase separation into droplets with tunable viscosity and dynamics. Proc Natl Acad Sci U S A 112, 7189–7194 (2015).

21. Posey, A.E., Holehouse, A.S. & Pappu, R.V. Phase Separation of Intrinsically Disordered Proteins. Methods Enzymol 611, 1–30 (2018).

22. Cahn, J.W. Phase separation by spinodal decomposition in isotropic systems. J Chem Phys 93, 93–99 (1965).

23. Sear, R.P. Phase separation of equilibrium polymers of proteins in living cells. Faraday Discuss 139, 21–34; discussion 105-128, 419-120 (2008).

24. Brangwynne, C.P., Tompa, P. & Pappu, R.V. Polymer physics of intracellular phase transitions. Nat Phys 11, 899–904 (2015).

25. Flory, P.J. Thermodynamics of high polymer solutions. J Chem Phys 51, 51–61 (1942).

26. Lin, Y.H., Forman-Kay, J.D. & Chan, H.S. Theories for Sequence-Dependent Phase Behaviors of Biomolecular Condensates. Biochemistry 57, 2499–2508 (2018).

27. Cates, M.E. & Tailleur, J. Motility-induced phase separation. Annu Rev Condens Matter Phys 6, 219–244 (2015).

28. Ranganathan, S. & Shakhnovich, E.I. Dynamic metastable long-living droplets formed by sticker-spacer proteins. Elife 9(2020).

29. Zhang, Z. & Glotzer, S.C. Self-assembly of patchy particles. Nano Lett 4, 1407–1413 (2004).

30. Witten Jr, T. & Sander, L.M. Diffusion-limited aggregation, a kinetic critical phenomenon. Physical review letters 47, 1400 (1981).

31. Chen, J. et al. Single-molecule dynamics of enhanceosome assembly in embryonic stem cells. Cell 156, 1274–1285 (2014).

32. Hansen, A.S., Amitai, A., Cattoglio, C., Tjian, R. & Darzacq, X. Guided nuclear exploration increases CTCF target search efficiency. Nat Chem Biol 16, 257–266 (2020).

33. Izeddin, I. et al. Single-molecule tracking in live cells reveals distinct target-search strategies of transcription factors in the nucleus. Elife 3(2014).

34. Normanno, D. et al. Probing the target search of DNA-binding proteins in mammalian cells using TetR as model searcher. Nat Commun 6, 7357 (2015).

35. Gautier, A. et al. An engineered protein tag for multiprotein labeling in living cells. Chem Biol 15, 128–136 (2008).

36. Grimm, J.B. et al. A general method to fine-tune fluorophores for live-cell and in vivo imaging. Nat Methods 14, 987–994 (2017).

37. Torreno-Pina, J.A. et al. Enhanced receptor-clathrin interactions induced by N-glycan-mediated membrane micropatterning. Proc Natl Acad Sci U S A 111, 11037–11042 (2014).

38. Wright, R.H. et al. ADP-ribose-derived nuclear ATP synthesis by NUDIX5 is required for chromatin remodeling. Science 352, 1221–1225 (2016).

39. Muñoz-Gil, G., Garcia-March, M.A., Manzo, C., Martin-Guerrero, J.D. & Lewenstein, M. Single trajectory characterization via machine learning. New J Phys 22, 013010 (2020).

40. Massignan, P. et al. Nonergodic subdiffusion from Brownian motion in an inhomogeneous medium. Phys Rev Lett 112, 150603 (2014).

41. Mandelbrot, B.B. & Van Ness, J.W. Fractional brownian motions, fractional noises and applications. SIAM Rev 10, 422–437 (1968).

42. Manzo, C. et al. Weak ergodicity breaking of receptor motion in living cells stemming from random diffusivity. Phys Rev X 5, 011021 (2015).

43. Guigas, G., Kalla, C. & Weiss, M. Probing the nanoscale viscoelasticity of intracellular fluids in living cells. Biophys J 93, 316–323 (2007).

44. Pageon, S.V., Nicovich, P.R., Mollazade, M., Tabarin, T. & Gaus, K. Clus-DoC: a combined cluster detection and colocalization analysis for single-molecule localization microscopy data. Mol Biol Cell 27, 3627–3636 (2016).

45. Berry, J., Weber, S.C., Vaidya, N., Haataja, M. & Brangwynne, C.P. RNA transcription modulates phase transition-driven nuclear body assembly. Proc Natl Acad Sci U S A 112, E5237–5245 (2015).

46. Szabo, F. The linear algebra survival guide:illustrated with Mathematica. Academic Press (2015).

47. Lee, D.S.W., Wingreen, N.S. & Brangwynne, C.P. Chromatin Mechanics Dictates Subdiffusion and Coarsening Dynamics of Embedded Condensates. bioRxiv, 2020.2006.2003.128561 (2020).

48. Garcia, D.A. et al. A New Model for Single-Molecule Tracking Analysis of Transcription Factor Dynamics. bioRxiv, 637355 (2019).

## SUPPLEMENTARY REFERENCES

1. Binder, K. Theory of first-order phase transitions. Reports on progress in physics 50, 783 (1987).

2. De Gennes, P.-G. & Gennes, P.-G. Scaling concepts in polymer physics. Cornell university press, 1979.

3. Gibbs, J.W. The Scientific Papers of J. Willard Gibbs: Dynamics. Dover Publ., 1961.

4. Debenedetti, P.G. Metastable liquids: concepts and principles, vol. 1. Princeton university press, 1996.

5. Oxtoby, D.W. Homogeneous nucleation: theory and experiment. Journal of Physics: Condensed Matter 4, 7627 (1992).

6. Pruppacher, H. & Klett, J. Microphysics of clouds and rainfall. Ed. Kluwer Academic Publishers Dordrecht, Netherlands (1997).

7. Sear, R.P. Nucleation: theory and applications to protein solutions and colloidal suspensions. Journal of Physics: Condensed Matter 19, 033101 (2007).

8. Witten Jr, T. & Sander, L.M. Diffusion-limited aggregation, a kinetic critical phenomenon. Physical review letters 47, 1400 (1981).

9. Kartha, M.J. & Sayeed, A. Phase transition in diffusion limited aggregation with patchy particles in two dimensions. Physics Letters A 380, 2791–2795 (2016).

10. Szabo, F. The linear algebra survival guide:illustrated with Mathematica. Academic Press (2015).

11. Karim, F., Majumdar, S., Darabi, H. & Chen, S. LTSM fully convolutional networks for time series classification. IEEE access 6, 1662–1669 (2017).

12. Scher, H. & Montroll, E.W. Anomalous transit-time dispersion in amorphous solids. Phys Rev B 12, 2455 (1975).

13. Mandelbrot, B.B. & Van Ness, J.W. Fractional brownian motions, fractional noises and applications. SIAM Rev 10, 422–437 (1968).

14. Massignan, P. et al. Nonergodic subdiffusion from Brownian motion in an inhomogeneous medium. Phys Rev Lett 112, 150603 (2014).

15. Lim, S. & Muniandy, S. Self-similar gaussian processes for modeling anomalous diffusion. Phys Rev E 66, 021114 (2002).

16. Von Der Ahe, D. et al. Glucocorticoid and progesterone receptors bind to the same sites in two hormonally regulated promoters. Nature 313, 706–709 (1985).

17. Presman, D.M. et al. DNA binding triggers tetramerization of the glucocorticoid receptor in live cells. Proceedings of the National Academy of Sciences 113, 8236–8241 (2016).

18. Tinevez, J.-Y. et al. TrackMate: An open and extensible platform for single-particle tracking. Methods 115, 80–90 (2017).

19. Ester, M., Kriegel, H.-P., Sander, J. & Xu, X. A density-based algorithm for discovering clusters in large spatial databases with noise. Kdd; 1996; 1996. p. 226–231.

20. Fujiwara, T., Ritchie, K., Murakoshi, H., Jacobson, K. & Kusumi, A. Phospholipids undergo hop diffusion in compartmentalized cell membrane. J Cell Biol 157, 1071–1081 (2002).

21. Sadegh, S., Higgins, J.L., Mannion, P.C., Tamkun, M.M. & Krapf, D. Plasma Membrane is Compartmentalized by a Self-Similar Cortical Actin Meshwork. Phys Rev X 7(2017).

22. Hansen, A.S., Amitai, A., Cattoglio, C., Tjian, R. & Darzacq, X. Guided nuclear exploration increases CTCF target search efficiency. Nat Chem Biol 16, 257–266 (2020).

